# Basicity of N5 in semiquinone enhances the rate of respiratory electron outflow in *Shewanella oneidensis* MR-1

**DOI:** 10.1101/686493

**Authors:** Yoshihide Tokunou, Keisuke Saito, Ryo Hasegawa, Kenneth H. Nealson, Kazuhito Hashimoto, Hiroshi Ishikita, Akihiro Okamoto

## Abstract

Extracellular electron transport (EET) occurs in environmental iron-reducing bacteria and is mediated by an outer membrane multi-heme cytochrome complex (Cyts). It has critical implications for global mineral cycling and electrochemical microbial catalysis. The rate of EET mediated by multiple heme redox centers significantly increases in the presence of flavins and quinones. Their electron free energy does not entirely account for the fact that differential effects on EET rate enhancement vary significantly by factors ≥100. Here, we report on whole-cell electrochemical analysis of *Shewanella oneidensis* MR-1 using six flavin analogs and four quinones. We demonstrated that protonation of the nitrogen atom at position 5 (N5) of the isoalloxazine ring is essential for electron outflow acceleration as a bound non-covalent cofactor of Cyts. EET mediated by Cyts was accelerated at a rate dependent on p*K*_a_(N5). The EET rate largely decreased in response to the addition of deuterated water (D_2_O), while low concentration of D_2_O (4 %) had little impact on electron free energy difference of the heme and non-covalent bound cofactors, strongly suggesting that the protonation of N5 limits the rate of EET. Our findings directly link EET kinetics to proton transport reaction via N5 and provide a basis for the development of novel strategies for controlling EET-associated biological reactions.

**Significance statement:** The potential of various small molecules such as flavins and quinones to enhance the rate of extracellular electron transport (EET) has been exploited to develop environmental energy conversion systems. Flavins and quinones have similar molecular structures but their abilities to enhance EET vary by >100× in *Shewanella oneidensis* MR-1. These large differences are inconsistent with conventional models, which rely on redox potentials or diffusion constant of shuttling electron mediators. In this study, we demonstrated that the basicity of the nitrogen atom of the isoalloxazine ring (N5) enhances the rate of electron outflow when a flavin or quinone is a non-covalent cofactor of *S. oneidensis* MR-1 outer membrane *c*-type cytochromes.

## Introduction

Extracellular electron transport (EET) is the process through which prokaryotes move electrons between the membrane interior and the cell exterior, such that redox reactions can occur with otherwise unavailable extracellular electron donors or acceptors (1-4). Microbes with the EET molecular machinery can render innovative metabolic processes inaccessible to microbes without the EET machinery. These include redox reactions with solid minerals (1, 2), intercellular electron transfer (5, 6), and surface interactions with electrodes for application in energy production and environmental technologies (7-11). The role of redox molecules as enhancers of bacterial current production remains controversial. High exogenous quinone concentrations (≥10 µM) enhance EET via an electron-shuttling mechanism, which is a two-electron redox process (12-14). In contrast, low levels of endogenous riboflavin or flavin mononucleotide (FMN) (≤1 µM) accumulate in cultures of EET-capable bacteria, non-covalently bind specific sites of outer membrane *c*-type cytochromes (Cyts) as cofactors (15-18), and form singly reduced semiquinone (SQ). Upon one-electron reduction of flavin, protonation occurs at the N5 nitrogen with the p*K*_a_ value (p*K*_a_(N5)) of 8.6 (19).

SQ enhances the EET rate to the same extent as exogenous quinone-mediated shuttling despite hundredfold lower endogenous concentrations (15, 18, 20-23). Nevertheless, electron transfer mediated by SQ are thermodynamically unfavorable. The redox potentials (*E*_*0*_) of the hemes in Cyts vary widely depending on the surrounding environment and are typically in the range of −400 to +200 mV relative to a standard hydrogen electrode (SHE) (24-32). SQ-mediated EET acceleration occurs when *E*_*0*_ for Cyts is >+50 mV (vs. SHE) (15, 18, 20). SQ and oxidized flavin cycling have *E*_*0*_ of ∼ −100 mV (vs. SHE), which is >150 mV lower than those of the Cyt heme groups (15, 18, 20). Therefore, thermodynamically unfavorable electron transfer from Cyts to SQ is unlikely to enhance the EET rate. Bound and free flavins are crucial in numerous anaerobic electron bifurcation reactions using them as SQ intermediates (33, 34). Flavin-based EET functions as a major metabolic pathway in various Gram-positive bacteria such as the pathogen *Listeria monocytogenes* (35, 36) and in other metal reducing bacteria *Geobacter sulfurreducens* PCA (16).

The interactions of flavins with purified Cyts have been extensively investigated in the model EET-capable bacterium *Shewanella oneidensis* MR-1 (17, 37-40). On the other hand, stabilization of the SQ state in Cyts was confirmed only in intact cells (15, 20) possibly because of the structural complexity and dynamics of Cyt (41-44). Whole-cell approaches significantly limit access to molecular-level information about specific enzymes. Consequently, SQ-mediated EET rate enhancement mechanisms are unknown. Recently, a whole-cell electrochemical assay demonstrated that molecules analogous to riboflavin can associate with Cyts in their single-electron reduced state (22). Thus, there is a potential control mechanism for the interaction between the Cyts binding site and the SQ cofactor.

In the present study, we analyzed the influence of (i) various flavin analogs and quinones on EET rate enhancement with respect to the p*K*_a_ of the flavin isoalloxazine ring; (ii) pH on the *E*_*0*_ for flavins and flavin analogs; and (iii) the kinetic isotope effect (KIE) on EET with heavy water using intact *S. oneidensis* MR-1 cells. These data indicate that the nitrogen atom at position-5 (N5) of the isoalloxazine ring determines the ability of the flavin analog to bind Cyts and forms a semi-reduced (SR) intermediate analogous to SQ before rapid and direct single-electron transfer. The protonation probability of N5 (i.e., p*K*_a_(N5)) is most likely a key determinant of the kinetic properties of EET.

## Results and discussion

To identify the key properties of the flavin analogs and quinones involved in EET kinetics, we measured the rate and amount of current produced by *S. oneidensis* MR-1 during lactate oxidation (*i*_*c*_) in the presence of riboflavin, flavin mononucleotide (FMN), six flavin analogs, and four quinones (Fig. 1A). We measured the formation of the semi-reduced (SR) state during *i*_*c*_ generation with each flavin and quinone using a three-electrode electrochemical system equipped with an indium tin-doped oxide (ITO) working electrode under +0.4 V (vs. SHE). Lactate (10 mM) served as the sole electron donor. Unless otherwise noted, the concentration of all flavin analogs and quinones tested was 2.0 μM. At this concentration, riboflavin strongly enhances *i*_*c*_ in *S. oneidensis* MR-1 as a bound cofactor by forming the intermediate SR state (15, 20). Flavin analogs with N5 in their polycyclic backbones (Fig. 1A) enhanced *i*_*c*_ at a rate equal to, or higher than that measured with riboflavin (Figs. 1B and S1A). MB gave the highest enhancement among N5 molecules. The dramatic *i*_*c*_ enhancement induced by MB was impaired by >90% via the deletion of the genes corresponding to Cyts (Fig. S1E). Similar substantial current decreases were also observed with riboflavin and FMN (15, 20). These data suggest that the N5-containing molecules enhance the rate of EET through Cyts, and their interactions with outer-membrane make negligible contributions to *i*_*c*_ as observed in several redox molecules (45, 46).

**Fig. 1.**
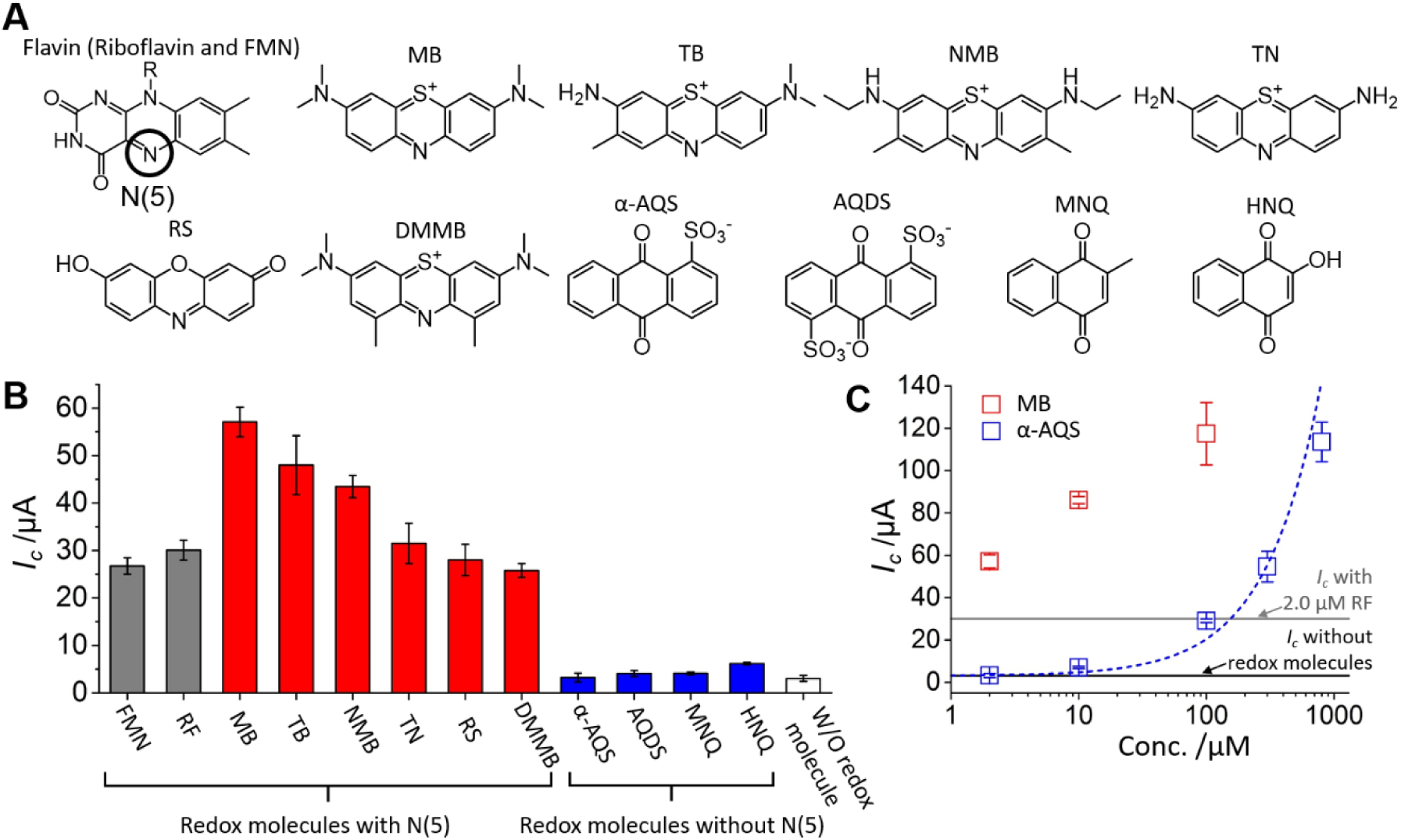
(A) Chemical structures of the redox molecules used in the present study. N5 in the isoalloxazine ring is circled in the chemical structure of flavin. Methylene blue (MB), toluidine blue (TB), new methylene blue (NMB), thionine (TN), resorufin (RS), and 1,9-dimethyl-methylene blue (DMMB) have N5, while anthraquinone-1-sulfonate (α-AQS), anthraquinone-1,5-disulfonate (AQDS), 2-methyl-1,4-naphthoquinone (MNQ), and 2-hydroxy-1,4-naphthoquinone (HNQ) lack it. (B) Maximum catalytic current of microbial lactate oxidation in *S. oneidensis* MR-1 (*I*_*c*_) after 10 h measurement in the presence of each molecule shown in (A). *I*_*c*_ in the presence of flavin, flavin analogs, and quinones are represented as gray, red, and blue bars, respectively. Concentration of each redox molecule was set to 2.0 µM. Error bars represent mean ± SEM for ≥three individual experiments in separate reactors. (C) *I*_*c*_ vs. MB or α-AQS concentration in the reactor. Blue dotted line represents *I*_*c*_ estimated by Fick’s law according to the diffusion kinetics of α-AQS between the cell and the electrode. Error bars represent the mean ± SEM for ≥three individual experiments in separate reactors.

Comparatively less EET enhancement was observed in the presence of quinones lacking N5 (blue bars in Figs. 1B and S1B). We compared the concentration dependencies of MB (with N5) and α-AQS (without N5) over 10 h using the max *i*_*c*_ value (*I*_*c*_). *I*_*c*_ increased with α-AQS concentration in accordance with the α-AQS diffusion kinetics estimated by Fick’s law (Figs. 1C and S1C). A concentration of 100 µM α-AQS was required to reach the same level of *i*_*c*_ as that achieved using 2.0 µM riboflavin, which is characteristic of diffusion-based shuttling mechanisms (Fig. 1C). The diffusion-limited kinetics of electron transfer from α-AQS to the electrode was previously confirmed by voltammetric analysis at scan rates of 1– 100 mVs^−1^ in the presence of the *S. oneidensis* MR-1 biofilm (47). With an N5-containing molecule (N5 molecule), MB, *I*_*c*_ enhancement did not follow Fick’s law and occurred at low concentrations (Fig. 1C and S1D). The observed scan rate dependency in cyclic voltammetry analysis indicated that the rate of electron transfer is not limited by diffusion kinetics in MB or riboflavin (Fig. S2). However, their kinetics were limited by diffusion in the absence of *S. oneidensis* MR-1 (Fig. S2). These data show that N5 molecules did not enhance *i*_*c*_ by a shuttling mechanism but rather by a direct electron transport process at the cell/electrode interface.

Stabilization of the N5 molecules in their SR form in Cyts was confirmed by the number of electrons involved in the redox reactions of all N5 molecules via differential pulse voltammetry (DPV) estimated from the half-width (Δ*E*_*p/2*_) of the oxidation peak (48, 49) (Figs. S3 and S4). Unbound MB in a cell-free system showed a Δ*E*_*p/2*_ of 55 mV which indicates a two-electron redox reaction (Fig. S3) (48, 49). The presence of *S. oneidensis* MR-1 cells changed the Δ*E*_*p/2*_ of MB to 120 mV (Fig. S3). This value resembles that reported for the one-electron redox reaction of flavin bound with Cyts (130 mV) (15, 20). Therefore, MB is stabilized in its SR form in the presence of MR-1. The peak MB current decreased upon the addition of the free radical scavenger α-tocopherol (Fig. S5), which supports the formation of SR state (15, 50). The oxidative signal of MB showed a single peak corresponding to the SR state up to a concentration of 10 µM (Fig. S6). Thereafter, the increase of MB concentration gradually shifted the peak potential (*E*_*p*_) towards the *E*_*p*_ of oxidation peak in two-electron reaction (Fig. S7). Therefore, MB would serve as a bound cofactor below 10 µM concentration in our experimental setup. In contrast, a one-electron oxidation peak was not detected for α-AQS (without N5) in the DP voltammogram (Fig. S8). These data demonstrate that MB enhances the rate of EET in the SR state. The same trend for Δ*E*_*p/2*_ was observed with TB, NMB, TN, RS, and DMMB (Fig. S4). These results suggest that the N5 atom in the polycyclic backbone is the critical structural component of flavin analogs, which enables them to function in the SR state as bound cofactors in Cyts. The formation of the SR state with electron flow enhancement is reminiscent of the SQ states observed in numerous microbial electron bifurcation reactions (33). Some of these have been confirmed in the crystal structures of flavodoxins in the oxidized quinone, semiquinone, and hydroquinone forms (51). It seems possible that the SR state is stabilized by donation of a hydrogen bond to N5 from the Cyts scaffold, as observed in flavodoxins (52).

To confirm whether the redox cycle of bound cofactors could couple with protonation as with that of SQ in flavodoxins, we evaluated the effects of bulk solution pH on the redox potential (*E*_*0*_) using the oxidation peaks in DP voltammograms. When a bulk solution was acidified in the presence of *S. oneidensis* MR-1 and each cofactor at 2 μM, the *E*_*0*_ of the N5 molecules, including riboflavin, linearly decreased upon pH increase, which indicates a proton-coupled electron transfer reaction (Fig. 2). On the other hand, the observed slope values were lower than expected (riboflavin: −13 mV pH^−1^; MB: −34 mV pH^-1^; TB: −32 mV pH^-1^; NMB: −30 mV pH^-1^; and TN: −21 mV pH^−1^ (Fig. 2)). Similar deviations from the −59 mV pH^-1^ slope were reported for SQs in certain flavodoxins and flavoproteins forming hydrogen bonds with protein scaffolds via N5 (53-55). The absence of the corresponding deviation in the free state flavins and quinones (53, 55) suggests that the low pH-dependence of *E*_*0*_ could be due to conformational changes of the protein structure, which originates from changes of the protonation states of the titratable sites.

**Fig. 2.**
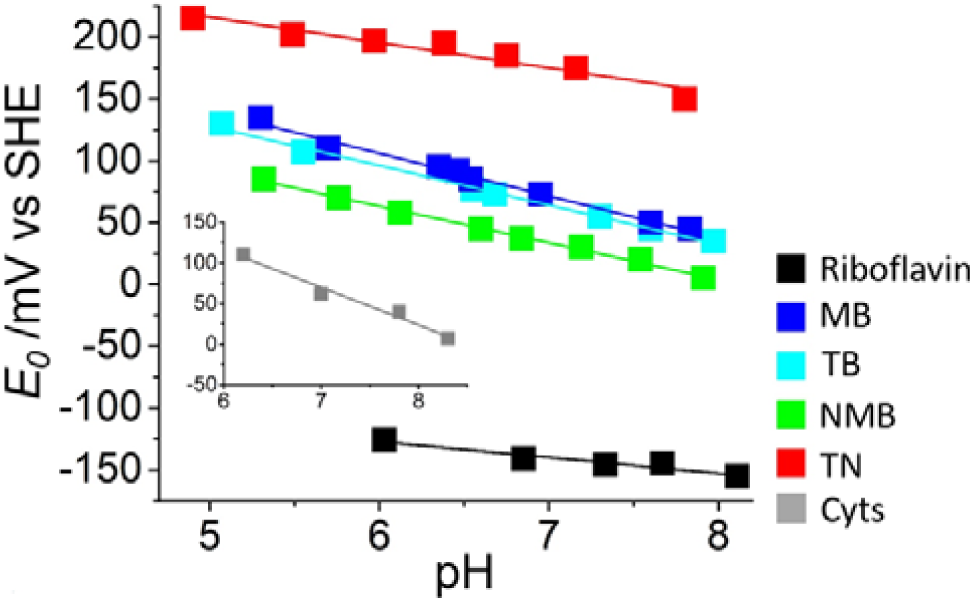
*E*_*0*_ of the bound cofactors (riboflavin, MB, TB, NMB, and TN) determined by DPV as a function of bulk solution pH. The slopes for the plots are riboflavin: −13 mV pH^−1^; MB: −34 mV pH^-1^; TB: −32 mV pH^−1^; NMB: −30 mV pH^-1^; and TN: −21 mV pH^−1^. Inset: *E*_*0*_ of Cyts derived from the DPV of riboflavin-bound Cyts. The slope was −47 mV pH^−1^ which is near the reported value for MtrC, a subunit of Cyts (56).

Given that the flavins receive electrons from the heme group(s) of Cyts, we previously posited that molecules with higher *E*_*0*_ could be more favorable for the acceleration of EET in our condition (+0.4 vs. SHE) (15, 32). It is evident from Fig. 1B that the extent of EET rate enhancement varies depending upon the binding N5 molecules. Unexpectedly, the EET rate did not increase with higher *E*_*0*_ in some N5 molecules (Fig. 3A). We approximated the capability of EET from the max *i*_*c*_ value (*I*_*c*_) in the 10-h measurements of each of the N5 molecules at the concentration of the Cyts complex with bound cofactors (Fig. S9 and Table S1). Their *E*_*0*_ were estimated by DPV (Figs. S3 and S4). It was assumed that the N5 molecules act as bound cofactors up to a concentration of 10 µM because of the DPV data (Figs. S6 and S7), and the EET capability for each Cyts forming a complex with bound cofactor was estimated from the dissociation constant (*K*_d_) at the N5 molecule concentrations of 2–10 µM (Fig. S9). As shown in Fig. 3A, the EET rate increased with *E*_*0*_ for the N5 molecules <+50 mV (vs. SHE). However, this tendency does not hold true for N5 molecules with higher *E*_*0*_. The EET rate for RS was low possibly due to a low overpotential to be oxidized by the electrode. Nevertheless, the cutoff was nearly identical to *E*_*0*_ for Cyts (+50 mV vs. SHE) (15, 32). The N5 molecules with *E*_*0*_ >+50 mV accept electrons from Cyts in a thermodynamically favorable downhill reaction. Thus, the suppression of EET rate could not originate from energetics of electron transfer.

**Fig. 3.**
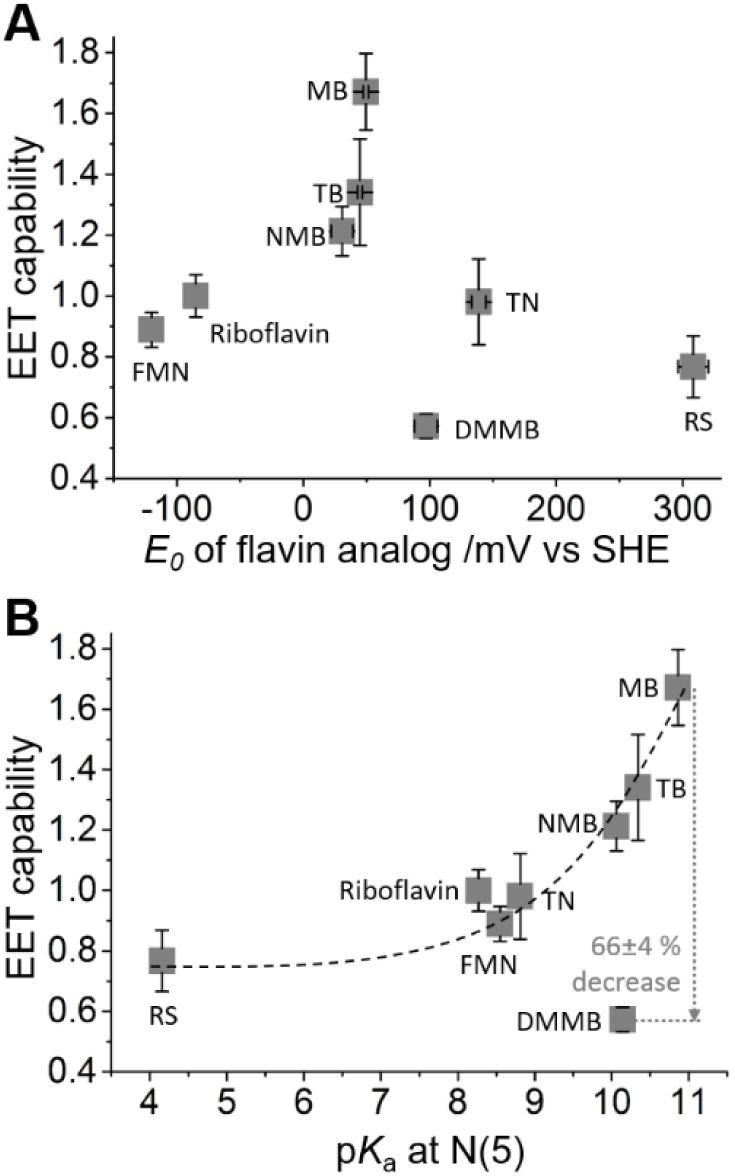
EET capabilities of *S. oneidensis* MR-1 with each N5 molecule as a function of the *E*_*0*_ of each cofactor (A) and p*K*_a_(N5) in one-electron reduced form (B). The *E*_*0*_ for all cofactors bound to Cyts except FMN and riboflavin were determined by differential pulse voltammetry (DPV) (Figs. S3 and S4). The *E*_*0*_ for FMN and riboflavin were obtained from the literature (15, 20). The p*K*_a_(N5) were calculated using a quantum chemical approach (Figs. S10 and S11). p*K*_a_(N5) of FMN and riboflavin were obtained from the literature (19). Error bars represent the mean ± SEM for ≥three individual experiments. The gray dotted line shows the suppression of the EET capability of DMMB relative to MB. The dashed line is for visual guidance and orientation.

We compared p*K*_a_(N5) with the EET capability of each N5 molecule. p*K*_a_(N5) in the SR state was calculated using a quantum chemical approach, as demonstrated for 28 reference compounds (Figs. S10 and S11) (57, 58). EET capability increased with p*K*_a_(N5) in the SR form (Fig. 3B). The increase in current production at high p*K*_a_(N5) suggests that the N5 protonation of the singly reduced cofactor limits the rate of EET. Nucleophilicity at N5 was also estimated by the Hammett substituent parameters. EET capability increased with the increase of nucleophilicity at N5 (Fig. S12 and Table S2). Meanwhile, DMMB with p*K*_a_ of 10.14 showed ∼66 ± 4% lower EET capability than MB (Fig 3B). Since DMMB has the identical backbone structure with MB except for methyl groups at the vicinity of N5 moiety in the isoalloxazine ring, strict suppression of EET capability in DMMB may be caused by prevention of protonation at N5. In contrast, there was no clear relationship between current production and p*K*_a_ value in the two-electron reduced form (Fig. S13). In the electron-shuttling mechanism involving a two-electron redox process, the rate limiting factors for EET are the diffusion constant and the *E*_*0*_ value (59). The distinct rate-limiting step from shuttling mechanism supports the bound-cofactor mechanism of the N5 molecules which mediate one-electron redox process as binding SR species, and supports the importance of proton uptake capability at N5 in EET kinetics.

To analyze the rate-limiting step in EET, we evaluated the kinetic isotope effect (KIE) using a highly reproducible *S. oneidensis* MR-1 monolayer biofilm (47, 60). We added deuterated water (≤ 4%; subtoxic concentration) to the bulk solution during current production of the MR-1 monolayer biofilm in the presence of 2.0 μM of each cofactor molecule. A previously reported experimental setup was used (47, 60). As the *i*_*c*_ was limited by the EET process, the KIE could characterize the rate-limiting proton transfer process through cofactor-bound Cyts (47). The influence of the KIE on the EET rate was evaluated from the differences in current production 10 min after D_2_O and H_2_O (*i*_*c*_(H_2_O)/*i*_*c*_(D_2_O)) were added to the electrochemical system. The amount of cofactor-bound Cyts complex was normalized by a dissociation constant (*K*_d_) in the presence and absence of 4% D_2_O (Fig. S9). D_2_O lowered current production within 10 s in the presence of TN (Fig. 4A). Further addition of D_2_O up to a final concentration of 4% continued to reduce the current (Fig. 4C). In contrast, the current increased by D_2_O addition in the presence of MB (Fig. 4B). The lower *K*_d_ value of MB was estimated in the presence of D_2_O (Fig. S9), and it caused the higher *i*_*c*_. The KIE for the cofactor-bound Cyts normalized to 1.06 ± 0.07, and those of riboflavin and TN were 1.35 ± 0.03 and 1.46 ± 0.10, respectively (Fig. 4D). These KIE values were larger than those measured in the presence of 100 μM α-AQS (gray dashed line in Fig. 4D), where the EET rate is limited by diffusion of α-AQS (47), strongly suggesting that the observed KIE values are assigned to riboflavin- and TN-bound Cyts. Therefore, the KIE data indicate that the EET mediated by cofactor-bound Cyts is limited by proton transport process. Substantial differences in KIE were detected when the cofactors in the Cyts were replaced. Therefore, each cofactor has its own unique KIE, and each N5 protonation/deprotonation most likely limits the EET rate. This finding is consistent with a previous report which suggested an association of rate-limiting proton transport with electron transport in flavin-bound Cyts (47).

**Fig. 4.**
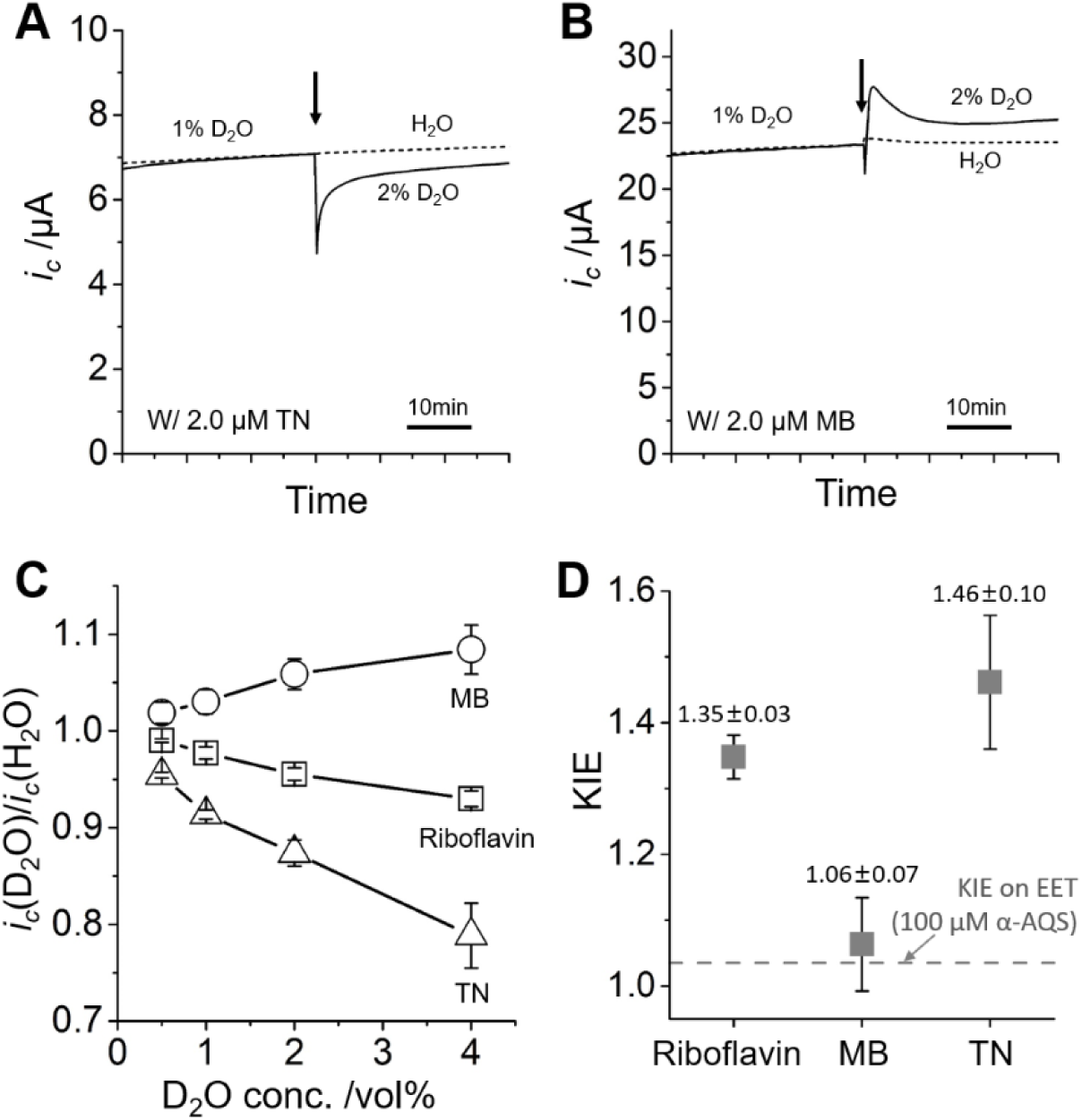
Effect of deuterium ion on EET kinetics in the presence of redox molecules. Representative time course for current production in the *S. oneidensis* MR-1 monolayer biofilm in the presence of 2.0 µM TN (A) and MB (B). The arrows indicate the time points of D_2_O (solid line) or H_2_O (dotted line) addition. Data corresponding to the dotted line were normalized to the data point immediately before D_2_O addition in the solid line data. (C) Effect of D_2_O addition at subtoxic concentrations (≤ 4% v/v) on *i*_*c*_ in the presence of 2.0 μM MB (circle plots) or TN (triangle plots). Data for 2.0 μM riboflavin (square plots) were obtained from the literature (47) for comparison. Error bars represent the mean ± SEM for ≥three experiments in separate reactors. (D) Kinetic isotope effect (KIE) on the EET per unit concentration of the cofactor-bound Cyts complexes in 4% (v/v) D_2_O. Error bars represent mean ± SEM for ≥three experiments in separate reactors. Gray dashed line represents the KIE on EET for 4% (v/v) D_2_O in the presence of 100 μM α-AQS reported in ref. (47). Numeric data are indicated above each plot.

The rate-limiting proton transfer is associated with the electron transfer from the hemes in the Cyts to the bound cofactors, and *E*_*0*_ potentially affect the EET rate (Fig. 3A). Consequently, any alteration in the *E*_*0*_ gap between Cyts and bound cofactors may increase the KIE values. To confirm that the observed KIE values were derived substantially from rate-limiting proton transfer rather than any change in *E*_*0*_ in the presence of D_2_O, we evaluated the *E*_*p*_ of Cyts, riboflavin, TN, and MB after D_2_O addition. Fig. 5 shows representative oxidation peaks obtained from the DP voltammograms of *S. oneidensis* MR-1 monolayer biofilm in the presence of 2.0 μM riboflavin, TN, or MB before and after the addition of 4% D_2_O. To determine *E*_*p*_ values, subtraction and deconvolution were conducted using an open source program SOAS (61). In D_2_O, not only a positive shift in the *E*_*p*_ of ∼25 mV for hemes in the Cyts but also a positive shift in the *E*_*p*_ of ∼25 mV for bound riboflavin were observed, without changing the *E*_*p*_ difference (Fig. 5A). The *E*_*p*_ for TN also positively shifted, and the *E*_*p*_ gap between Cyts and TN was also almost identical (Fig. 5B). Peaks of Cyts and MB were overlapped in both before and after addition of D_2_O (Fig. 5C), indicating little impact of D_2_O on *E*_*p*_ gap as well. Decrease of peak width in the presence of D_2_O may originate from the increase of MB/Cyts ratio contributing on the oxidation peak, which is consistent with higher affinity of MB with Cyts in the presence of D_2_O (Fig. S9). Collectively, *E*_*p*_ gap between Cyts and bound cofactors were scarcely influenced by D_2_O, further supporting that the observed KIE values in Fig. 4 were derived substantially from rate-limiting proton transfer via N5 of the bound cofactors.

The extent of current suppression by deuterated water was much larger than those determined when the EET kinetics is limited by diffusion process of proton donor for N5 (Fig. 4D). Assuming protons are delivered to N5 via diffusing water or buffer molecules in bulk solution and it limits the rate of EET, the ratio of the cofactor proceeding EET with D^+^/H^+^ is correlated with the percentage of heavy water added, i.e., the extent of current suppression is lower than the percentage of deuterated water added (62, 63). However, we observed the KIE values >1.04 in the presence of 4% D_2_O (Fig. 4D), indicating that diffusion of free water or buffer molecule do not limit the rate of EET. Some enzymes with several protonic sites transporting more than one proton suppress the rate of coupled electron transport over the ratio of added deuterated water as indicated in the Gross-Butler model (62, 64, 65). Given such protonation kinetics is not possible for free flavin or quinone, the observed large KIE further confirms our finding that the protonation of bound-cofactor limits the rate of EET via Cyts.

**Fig. 5.**
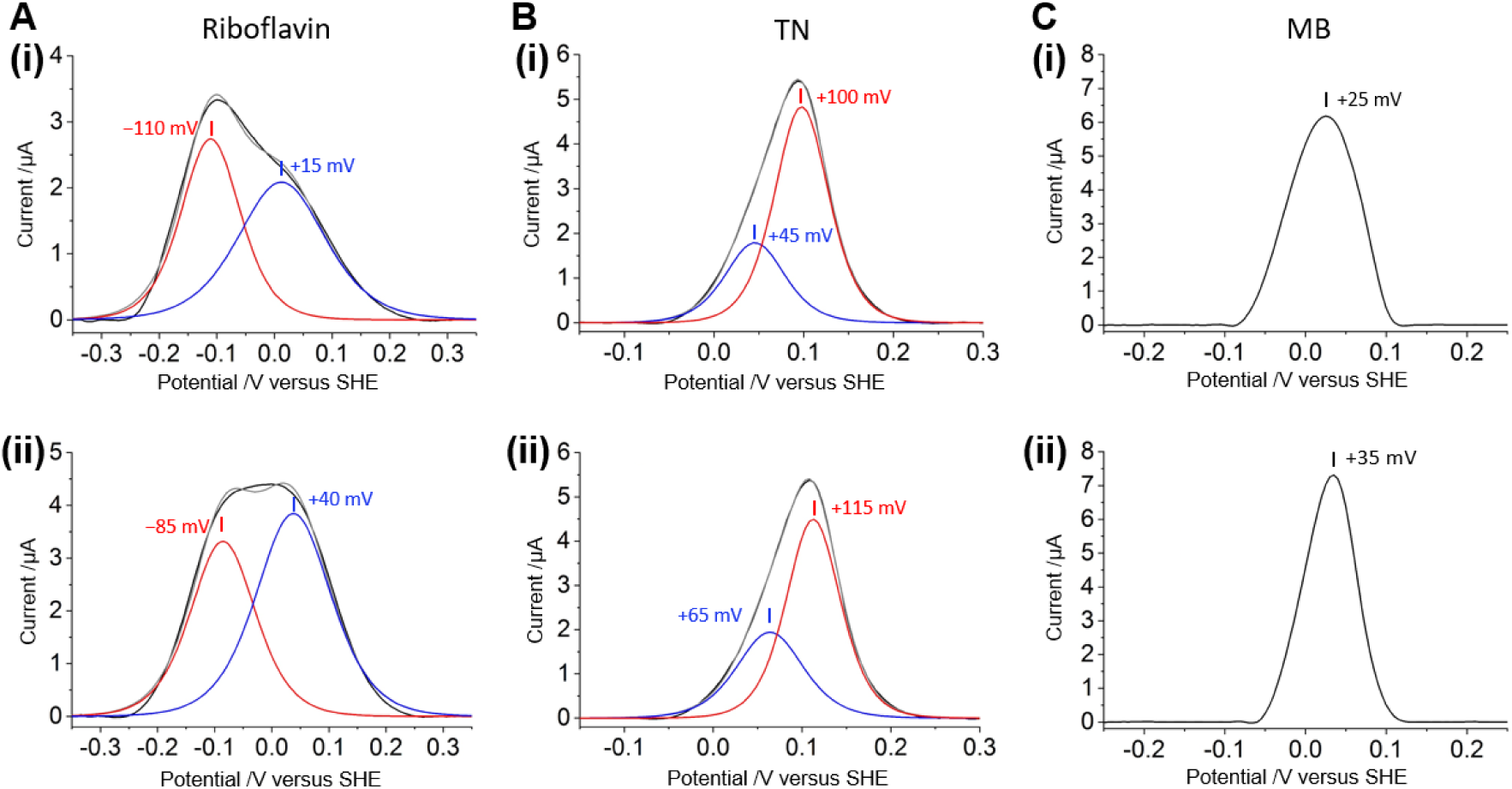
Baseline subtracted differential pulse (DP) voltammograms of the *S. oneidensis* MR-1 monolayer biofilm in the presence of 2.0 μM riboflavin (A), TN (B), and MB (C). (i) and (ii) represent the data before and after the addition of 4% (v/v) deuterium oxide, respectively. Black and gray lines are baseline subtracted DP voltammograms and fitted lines, respectively. A fitted line composed of two peaks is represented by red and blue lines. According to (15, 20), the blue peak is assigned to the Cyts and the red peak is assigned to a bound cofactor. The oxidative peaks in panel (C) were not able to be deconvoluted by SOAS because they overlapped.

The EET kinetics through Cyts was reported to be limited by proton transfer reaction in the absence of SR cofactor molecules (47). Therefore, high EET acceleration by cofactors may be the result of an increase in the rate of proton transfer coupled with the EET. Given EET links with the localization of protons across the inner-membrane as well in the absence of the bound cofactors (47, 66), it is of great interest to clarify detailed molecular mechanisms of EET-coupled complex proton transport to understand energy acquisition machinery of EET-capable bacteria in the presence and absence of the bound cofactors.

## Conclusions

We demonstrated that the basicity of N5 enhances the rate of EET when flavin is stabilized as SQ in Cyts of *S. oneidensis* MR-1. Therefore, high EET acceleration by the SQ intermediate in spite of unfavorable electron energetics may result from enhancement of proton transfer rate coupled with the EET. Significant variations in EET enhancement caused by flavin analogs and quinones are also explained by the function of N5, the proton uptake capability and the ability to form SR state binding with Cyts (Fig. 6). It is of great interest to test the function of N5 with the EET-capable bacterium *Geobacter sulfurreducens* and the pathogen *Listeria monocytogene* which use bound-flavin based EET mechanism as well (16, 35, 36). Understanding the role of the N5 and SR state associated with EET may help elucidate the function of redox bifurcation in biological systems, and could provide novel direction to control environmental and pathogenic bacterial activity.

**Fig. 6.**
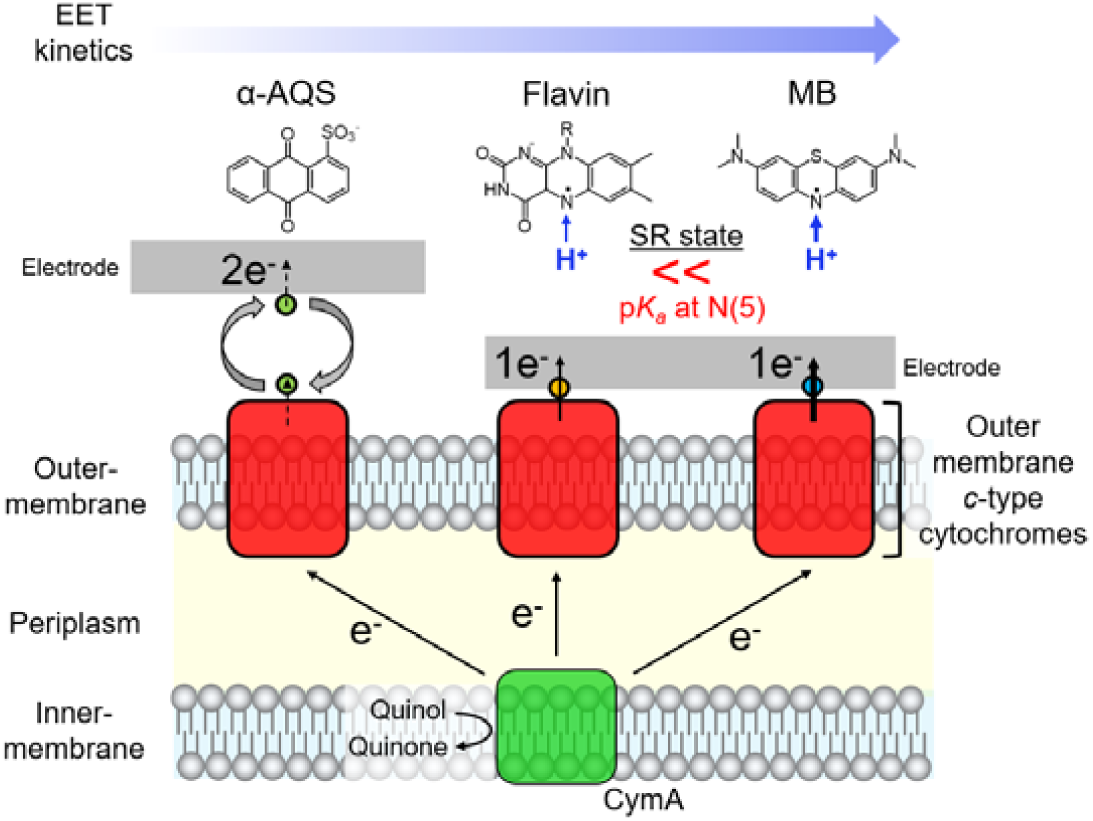
Schematic representation of the molecular control of respiratory electron outflow in *S. oneidensis* MR-1 mediated by flavin, flavin analogs, and quinones.

## Methods

### Strains and culture conditions

*Shewanella oneidensis* MR-1 cells were grown aerobically at 30 °C for 24 h in 15 mL Luria-Bertani (LB) medium (25 g L^-1^). The cell suspension was centrifuged at 6,000 × *g* for 10 min. The cell pellet was resuspended in 15 mL of a medium consisting of DM: NaHCO_3_ [2.5 g L^-1^], CaCl_2_·2H_2_O [0.08 g L^-1^], NH_4_Cl [1.0 g L^-1^], MgCl_2_·6H_2_O [0.2 g L^-1^], NaCl [10 g L^-1^], yeast extract [0.5 g L^-1^], and (2-[4-(2-hydroxyethyl)-1-piperazinyl] ethanesulfonic acid [HEPES; 7.2 g L^-1^] (pH 7.8) supplemented with 10 mM lactate as the carbon source for *S. oneidensis* MR-1. The cells were cultured aerobically at 30 °C for 12 h and centrifuged at 6,000 × *g* for 10 min. The cell pellet was washed twice with DM medium by centrifugation for 10 min at 6,000 × *g*. A mutant strain lacking the genes encoding Cyts (Δ*omcAll*; deletions of *SO1778-SO1782, SO2931*, and *SO1659*) was constructed as previously described (67).

### *Electrochemical measurements of the catalytic current generated by lactate oxidation (i*_*c*_*) in* S. oneidensis *MR-1 with each redox molecule*

A single-chamber, three-electrode system for whole-cell electrochemistry was constructed as previously described (15, 22). An indium tin-doped oxide (ITO) substrate (surface area: 3.1 cm^2^) was placed at the bottom of the reactor and used as the working electrode. Ag/AgCl (saturated KCl) and a platinum wire (surface area: ∼10 mm^2^) were used as the reference- and counter-electrodes, respectively. DM 4.0 mL (pH 7.8) containing each redox molecule (Fig. 1A) and 10 mM lactate as the sole electron donor was de-aerated by bubbling with N_2_ for > 20 min. It was then added to the electrochemical cell as an electrolyte. The concentration of flavin analogs and quinones was set to 2.0 μM unless otherwise indicated. The reactor was maintained at a temperature of 30 °C and was not stirred during measurements. Cell suspensions with OD_600_ = 0.1 were inoculated into the reactor. The working electrode was poised at +0.4 V vs. the standard hydrogen electrode (SHE). Electrochemistry-based experimental details about estimation of dissociation constant (*K*_d_), EET capability, and KIE value are described in Supporting information.

### *Formation of the* S. oneidensis *MR-1 monolayer biofilm on the ITO electrode*

DM 4.0 mL (pH 7.8) with 10 mM lactate was added to the electrochemical cell as an electrolyte and was de-aerated by bubbling with N_2_ for > 20 min. The *S. oneidensis* MR-1 cell suspension with OD_600_ = 0.1 was grown in the reactor. The working electrode was poised at +0.4 V vs. SHE in the presence of 10 mM lactate as the sole electron donor. The bacteria were incubated at 30 °C with no agitation for 25 h. Formation of the monolayer biofilm was confirmed by *in situ* confocal fluorescence microscopy as previously described (32).

### Voltammetry conditions

Cyclic voltammetry (CV) and di□erential pulse voltammetry (DPV) were conducted to determine the electrochemical properties of the flavin analogs and quinones and the *S. oneidensis* MR-1 monolayer biofilm via an automatic polarization system (VMP3; BioLogic Science Instruments, Seyssinet-Pariset, France). DPV was conducted under the following conditions: 5.0 mV pulse increments, 50 mV pulse amplitude (Δ*E*_*pa*_), 300 ms pulse width, and 5.0 s pulse period. *E*_*0*_ was approximated from the equation *E*_*0*_ = *E*_*p*_ + (Δ*E*_*pa*_/2) (48), thus, *E*_*0*_ was estimated to be 25 mV more positive than the peak potential (*E*_*p*_) observed in DPV. To determine the peak potential (*E*_*p*_), half-width (Δ*E*_*p/2*_), and oxidation peak intensity in DPV, the background current was subtracted by fitting the baseline of regions remote from the peak and assuming a similar smooth charging current throughout the peak region. The subtraction was performed in the open source program SOAS (61).

### *Calculation of the acid-base equilibrium dissociation constants (p*K_a_*) in redox active molecules*

To compute the absolute p*K*_a_ values, we employed a quantum chemical approach (57). In the deprotonation reaction of the protonated state (AH) to deprotonated state (A^−^) in aqueous solution, p*K*_a_ is defined as

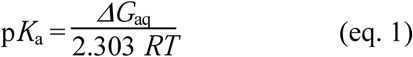

where *Δ G*_aq_ is the free energy difference between AH and (A^−^ + H^+^) in water (i.e., *Δ G*_aq_ = *G*_aq_(A^−^) + *G*_aq_(H^+^) –*G*_aq_(AH)), *R* is the gas constant, and *T* is the temperature. *Δ G*_aq_ can also be written as

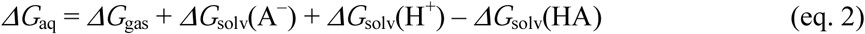

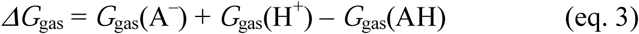

The free energy *G*_gas_ in vacuum can be obtained, using the following equation;

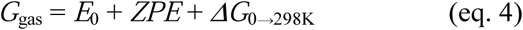

where *E*_0_ is the ground-state energy in vacuum, *ZPE* is the zero-point vibrational energy, and *Δ G*_0→298K_ is the thermal vibrational free energy at 298 K. For proton, the free energy *G*_gas_(H^+^) of 6.28 kcal/mol and *Δ G*_solv_(H^+^) of 265.74 kcal/mol were used (57). To obtain *E*_0_, *ZPE*, and *ΔG*_0→298K_, full geometry optimizations were carried out using the restricted DFT method for the non-radical states and the unrestricted DFT method for the radical states with the B3LYP functional and 6-31g** basis sets, and we used the Jaguar program code (68). Vibrational frequencies and electrostatic potentials were calculated using the geometry-optimized structures at the same level of theory. To calculate the ground-state electronic energy *E*_0_, we employed cc-pvqz basis sets for accuracy.

The solvation energy *ΔG*_solv_ was calculated by solving the Poisson equation using the Solvate module from MEAD (69), where van der Waals radii for H, N, O, Cl, and titratable H^+^ are 1.2, 1.4, 1.4, 1.9, and 1.0 Å, respectively; C for -CH_3_ and -CH_2_-groups are 2.0Å, whereas C for others are 1.2 Å (57). Atomic partial charges used for *ΔG*_solv_ were determined by the restraint-electrostatic-potential (RESP) method (70-72).

To evaluate the accuracy of the method, we calculated p*K*_a_ values for 28 compounds whose experimentally measured p*K*_a_ values are reported (i.e., ref. (73) for H_2_O and NH_4_^+^ and ref. (57) for other compounds) (Fig. S10B). We reproduced the experimentally measured p*K*_a_ values with a root-mean square deviation of 0.94 and a maximum error of 1.77 in p*K*_a_ units (Fig. S11).

## Acknowledgements

The authors thank Prof. Hiroyuki Noji and Prof. Kohei Uosaki for their helpful advice. This work was financially supported by JST CREST (No. JPMJCR1656), JSPS KAKENHI (No. 24000010 to K.H., No. 17H04969 to A. O., No. 17J02602 to Y.T., No. JP26800224 to K.S., No. JP16H06560 to K.S and H.I., and No. JP26105012 to H.I.), the Japan Agency for Medical Research and Development (AMED), the US Office of Naval Research Global (No. N62909-17-1-2038), the Materials Integration for Engineering polymers of the Cross-ministerial Strategic Innovation Promotion Program (SIP), and the Interdisciplinary Computational Science Program in CCS, University of Tsukuba. Y.T. is a JSPS Research Fellow and supported by JSPS through the Program for Leading Graduate Schools (MERIT).

## Author contributions

Y.T. and A.O. conceived and designed the study. Y.T. conducted the experiments and analyses. K.S., R.H., and H.I. conducted the theoretical calculations. Y.T., K.H.N., K.H., H.I., and A.O. wrote the manuscript.

