## Supporting Information for "Basicity of N5 in semiquinone enhances the rate of respiratory electron outflow in *Shewanella oneidensis* MR-1"

**SUPPLEMENTARY METHODS**

**Estimation of dissociation constant (*K*_d_) of complex formation between outer membrane *c*-type cytochromes (Cyts) and cofactor**

We estimated the dissociation constant (*K*_d_) of complex formation between Cyts and each N5 molecule, using the following equation.

$K_{d}= \frac{\left[ P \right]\left[ L \right]}{[PL]}$	(eq. S1)

where [P] is the concentration of Cyts, [L] is the concentration of unbound N5 molecule in solution, and [PL] is the concentration of the Cyts complex with the N5 molecule. *K*_d_ was estimated from the oxidation peak current in differential pulse voltammograms (DPV) before and after the addition of each N5 molecule, as previously described.^1^ Since it was assumed that the N5 molecules act as bound cofactors up to a concentration of 10 µM (Figures S6 and S7), *K*_d_ was estimated at the N5 molecule concentrations of 2–10 µM. When the peak intensity increased about α times with increase of the concentration of redox molecules from [L]_1_ to [L]_2_ and assumed that the sum of [P] and [PL] is constant throughout the measurement, *K*_d_ can be described as below, ^1^

$\alpha-1=K_{d}\left( \frac{1}{\left[ L \right]_{1}}-\frac{\alpha}{\left[ L \right]_{2}} \right)$	(eq. S2)

We estimated the *K*_d_ of complex formation between Cyts and each N5 molecule via linear regressions obtained from the plot of $\alpha-1$ against $\frac{1}{\left[ L \right]_{1}}-\frac{\alpha}{\left[ L \right]_{2}}$.

**Quantification of EET capability of *S. oneidensis* MR-1 in the presence of each redox molecule**

We approximated EET capability of *S. oneidensis* MR-1 with each N5 molecule using the maximum *i_c_* value (*I_c_*) during 10 h measurements, normalized by the amount of cofactor-bound Cyts complex. When the produced current is proportionally related to the amount of binding cofactor, the rate of EET per unit concentration of complex of Cyts and cofactor, β, can be described as follows.

$\beta= \frac{I_{c}}{[PL]}$	(eq. S3)

where [PL] represents the concentration of the Cyts bound with cofactor molecule. Incorporation of *K*_d_ provides:

$\beta=\frac{\left( [L]+K_{d} \right)I_{c}}{[L](\left[ P \right]+\left[ \mathrm{PL} \right])}$	(eq. S4)

In order to compare β among each N5 molecule, [L] was set as 2.0 μM and the sum of [P] and [PL] was assumed to be the same among each reactor. Normalization by the β of riboflavin (β_Riboflavin_) provides EET capability of each N5 molecule shown below.

$$EET capability :\frac{\beta}{\beta_{Riboflavin}}$$

**Addition of deuterated water (D_2_O) to the electrochemical reactor in the presence of a monolayer biofilm of *S. oneidensis* MR-1**

After formation of a monolayer biofilm of *S. oneidensis* MR-1 on an ITO electrode, supernatant solution was refreshed with anaerobic DM with 10 mM lactate containing either 2.0 µM methylene blue (MB) or 2.0 µM thionine (TN) at pH 7.8. Then D_2_O was sequentially added to the reactor at final concentrations ranging from 0.5 % to 4% (v/v).

**Normalization of the kinetic isotope effect (KIE) on EET using the amount of cofactor-bound Cyts complex**

The extent of kinetic isotope effect (KIE) on EET was evaluated from the difference between the current production 10 minutes after the addition of 4 % (v/v) D_2_O and H_2_O (*i_c_*(H_2_O)/*i_c_*(D_2_O)) to the electrochemical system in the presence of a monolayer biofilm of *S. oneidensis* MR-1, followed by normalization of the amount of cofactor-bound Cyts complex using dissociation constant (*K*_d_) in the presence of 4% D_2_O and in the absence of D_2_O as below:

$KIE=\frac{i_{c}(H_{2}O)}{i_{c}(D_{2}O)}\cdot\frac{{[PL]}_{D}}{{[PL]}_{H}}$ (eq. S5)

where [PL]_D_ and [PL]_H_ show the concentration of the Cyts bound with cofactor in the presence of 4% (v/v) D_2_O and in the absence of D_2_O, respectively. [PL] can be described from the definition of dissociation constant (*K*_d_) (eq. S1) as follows.

$[PL]=\frac{{[P]}_{\mathrm{total}}[L]}{K_{d}+[L]}$	(eq. S6)

where [P]_total_ is the sum of the concentration of Cyts unbound with cofactor and Cyts bound with cofactor, [L] is the concentration of unbound redox molecule in solution. Assuming that [P]_total_ is constant before and after the addition of D_2_O, the KIE can be described as follows.

$KIE=\frac{i_{c}(H_{2}O)}{i_{c}(D_{2}O)}\cdot\frac{K_{d}(H_{2}O)+[L]}{K_{d}(D_{2}O)+[L]}$	(eq. S7)

*K*_d_(D_2_O) and *K*_d_(H_2_O) represent the dissociation constant in the presence of 4% (v/v) D_2_O and in the absence of D_2_O, respectively. [L] was set as 2.0 μM.

**SUPPLEMENTARY DATA**

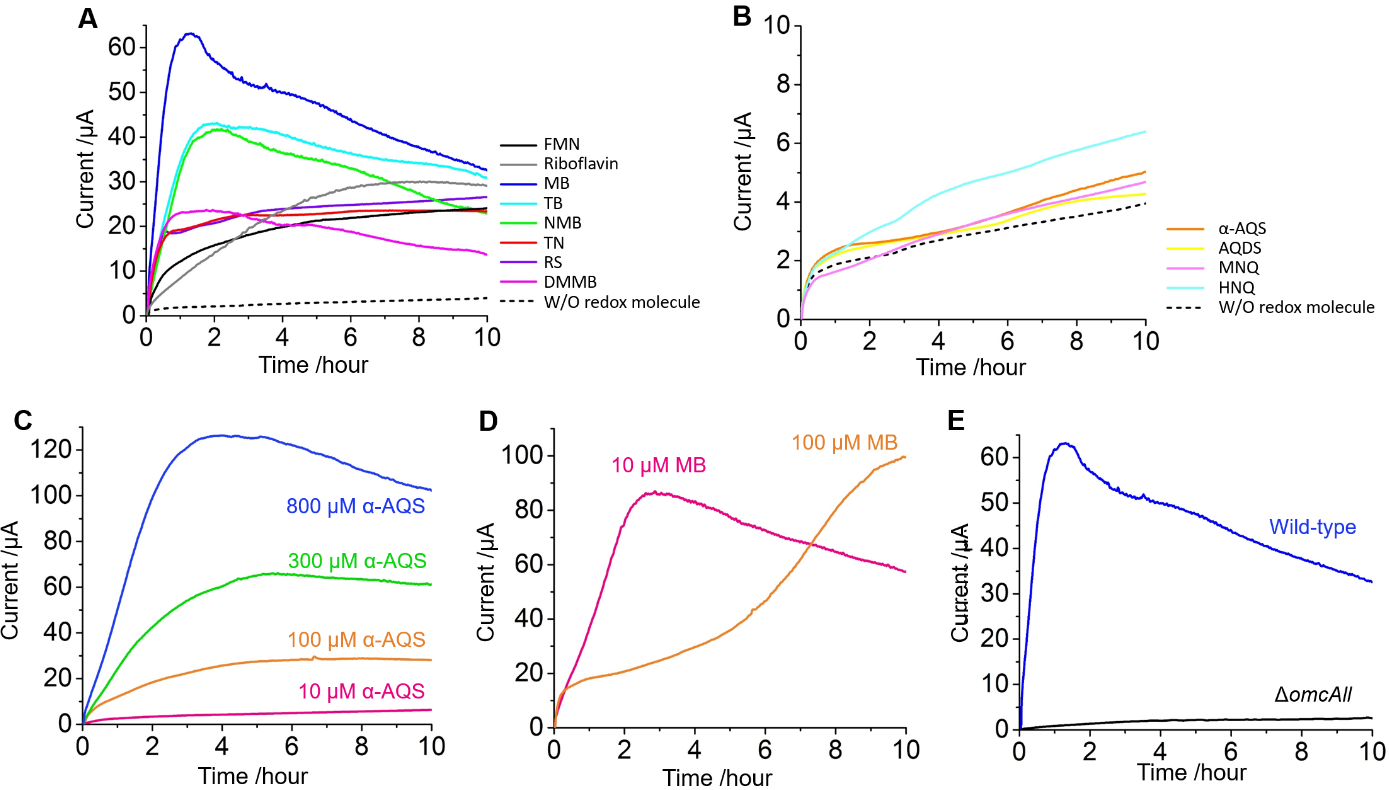

**Figure S1.** Current production from *S. oneidensis* MR-1. (**A**)-(**D**) Representative time courses of current generation from *S. oneidensis* MR-1 wild-type in the presence of 10 mM lactate on ITO electrode at +0.4 V (vs SHE). A cell suspension of *S. oneidensis* MR-1 in DM with an optical density of 0.1 at 600 nm (OD_600_ = 0.1) was inoculated at t = 0 in an electrochemical reactor containing each redox molecule shown in Figure 1A (solid lines) or without redox molecules (dotted line). The concentration of redox molecules was set as 2.0 μM unless noted. The same tendency was reproduced for each redox compound in at least three separate experiments. (**E**) Time course of current generation from *S. oneidensis* MR-1 wild-type (blue line) and gene deletion mutant corresponding with Cyts (Δ*omcAll*, black line) in the presence of 2.0 μM MB. The data of wild-type is the same with the blue line in Figure S1A.

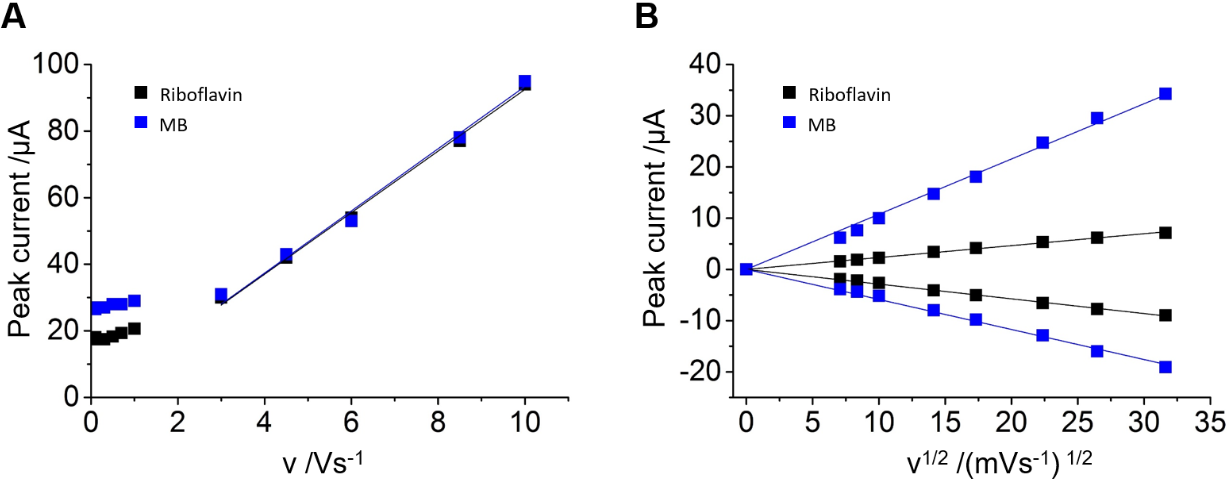

**Figure S2.** (**A**) Plots of peak current of 4.0 μM riboflavin (black plots) or 4.0 μM MB (blue plots) obtained in cyclic voltammetry (CV) in the presence of a monolayer biofilm of *S. oneidensis* MR-1 as a function of a scan rate. Both of peak current values linearly increased with the scan rate except for those when the metabolic current may affect the peak current due to slow scan rate. The linear relationships demonstrate that both riboflavin and MB are adsorbed and localized at the interface between cell membrane and electrode.^2^ (**B**) Plots of peak current of 4.0 μM riboflavin (black plots) or 4.0 μM MB (blue plots) in the absence of a monolayer biofilm of *S. oneidensis* MR-1 as a function of the square root of a scan rate. Those peak current values linearly increased with the square root of a scan rate, demonstrating that both riboflavin and MB proceeds redox reaction as diffusing species without a monolayer biofilm of *S. oneidensis* MR-1.

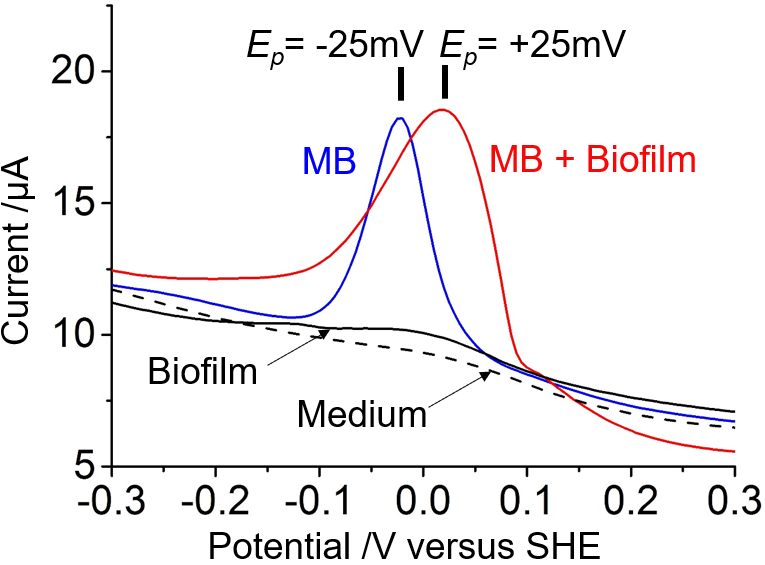

**Figure S3.** Differential pulse voltammograms (DPV) of MB. Red and blue lines represent the data of DPV of 2.0 μM MB with a monolayer biofilm of *S. oneidensis* MR-1 and without a monolayer biofilm of *S. oneidensis* MR-1, respectively. Black dashed line and solid line are DPV of DM medium with 10 mM lactate (DM-L) and that of DM-L with a monolayer biofilm of *S. oneidensis* MR-1, respectively. The half width (*ΔE_p/2_*) and the peak potential (*E_p_*) are determined after the background subtraction using an open source program SOAS.^3^

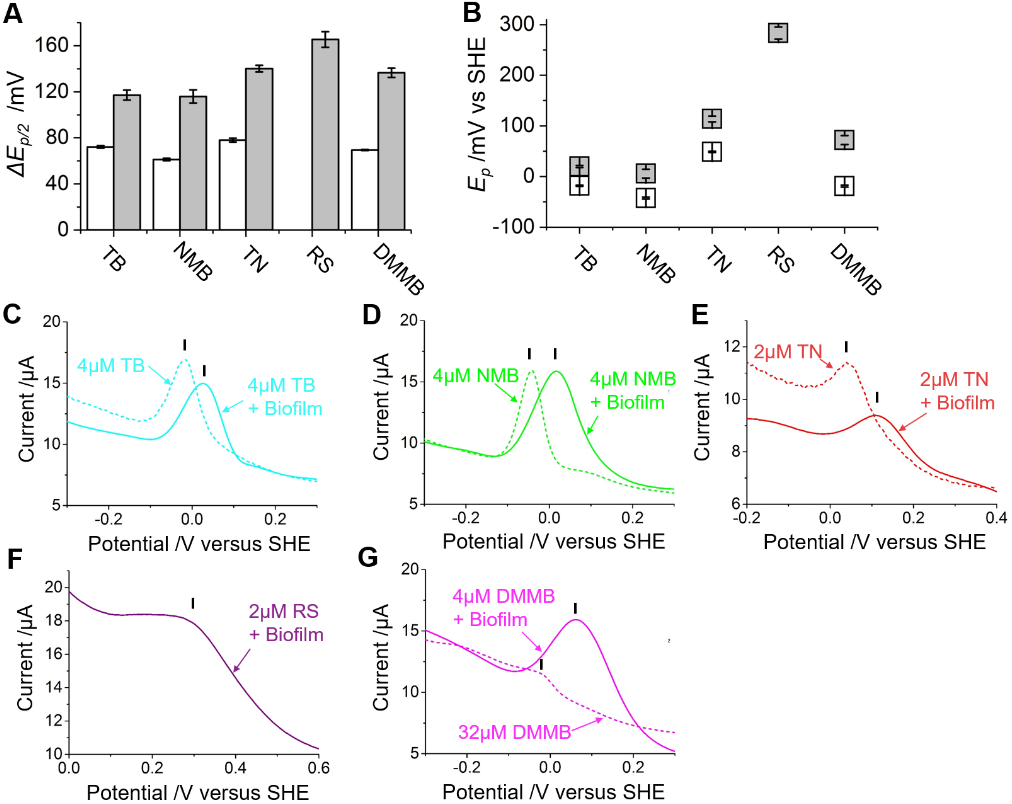

**Figure S4.** DPV of N5 molecules. The half width (*ΔE_p/2_*) (**A**) and the peak potential (*E_p_*) (**B**) of the oxidation peaks of N5 molecules in cell-free DM-L medium (white bars and plots) and those in the presence of a monolayer biofilm of *S. oneidensis* MR-1 (light gray bars and plots). The oxidation peak of RS in the cell-free condition was not detected. The standard errors in the *ΔE_p/2_* and *E_p_* are obtained from at least three individual DPV measurements in separate reactors. The representative DPV data of TB (**C**), NMB (**D**), TN (**E**), RS (**F**), DMMB (**G**) in the presence (solid lines) and absence (dashed lines) of a monolayer biofilm of *S. oneidensis* MR-1. The concentration of each molecule and the position of the oxidation peak are indicated.

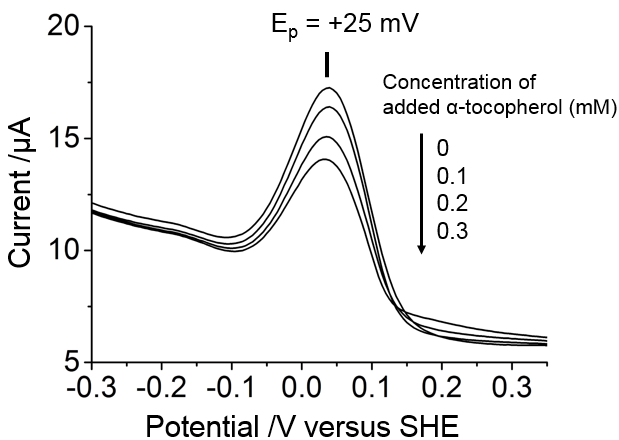

**Figure S5.** Effect of α-tocopherol addition on the DPV of 2.0 μM MB in the presence of a monolayer biofilm of *S. oneidensis* MR-1 on an ITO electrode. The concentrations of α-tocopherol added into the reactor are indicated.

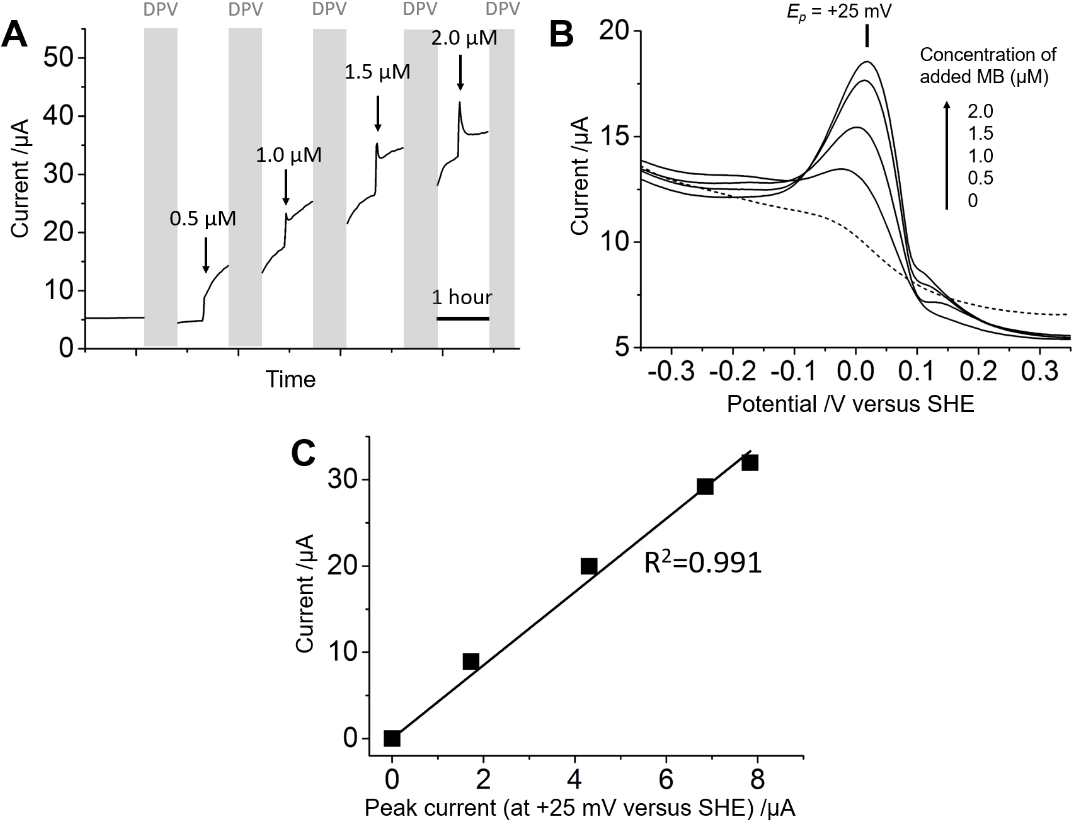

**Figure S6.** (**A**) Time course of a current production from *S. oneidensis* MR-1. The arrows and the gray regions indicate the timings of MB addition and DPV measurement, respectively. (**B**) DPV of *S. oneidensis* MR-1 containing 0.5, 1.0, 1.5, 2.0 μM MB (solid lines) and that in the absence of MB (dotted line), which were obtained during the measurement in (A). (**C**) Plots of the current production from a monolayer biofilm of *S. oneidensis* MR-1 against the oxidation peak current of MB at +25 mV versus SHE in DPV. Because the quantity of MB located at the cell-electrode interface correlated with the oxidation peak current at +25 mV versus SHE, this result indicates that the electrons produced from lactate oxidation in *S. oneidensis* MR-1 cells are predominantly delivered via the one-electron redox reaction of MB. The values of current production and peak current were obtained after subtraction of those in the absence of MB. The square of the correlation coefficients, R^2^ = 0.991, includes the point of origin.

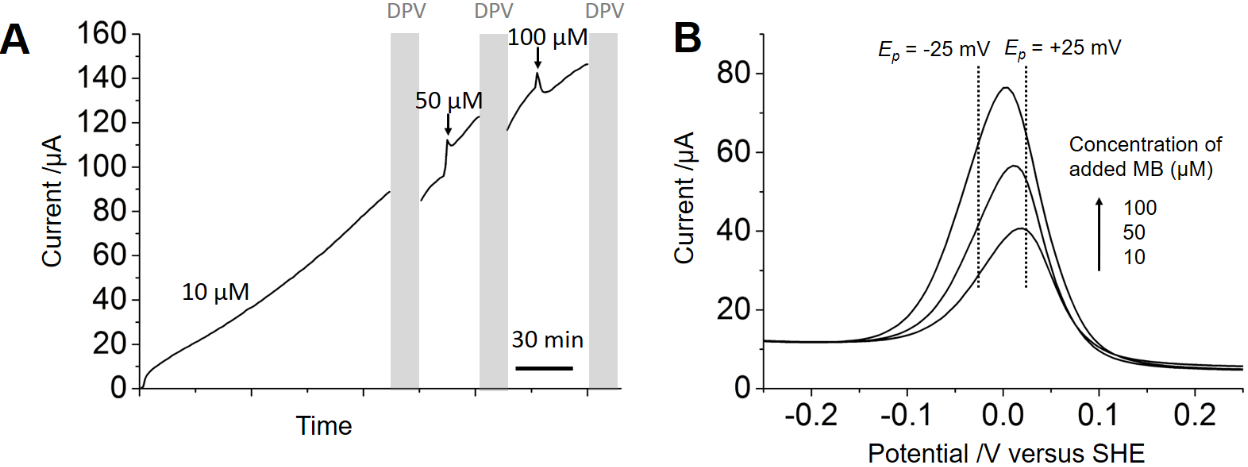

**Figure S7.** (**A**) Time course of a current production from *S. oneidensis* MR-1. The arrows and the gray regions indicate the timings of MB addition and DPV measurement, respectively. (**B**) DPV of *S. oneidensis* MR-1 containing 10, 50, 100 μM MB, which were obtained during the measurement in (A).

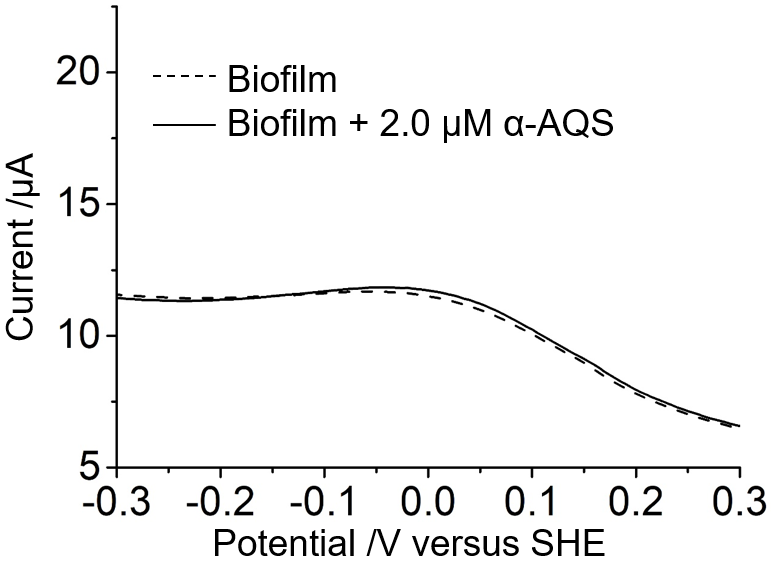

**Figure S8.** Representative DPV data of a monolayer biofilm of *S. oneidensis* MR-1 in the absence (dotted line) and presence (solid line) of 2.0 μM α-AQS.

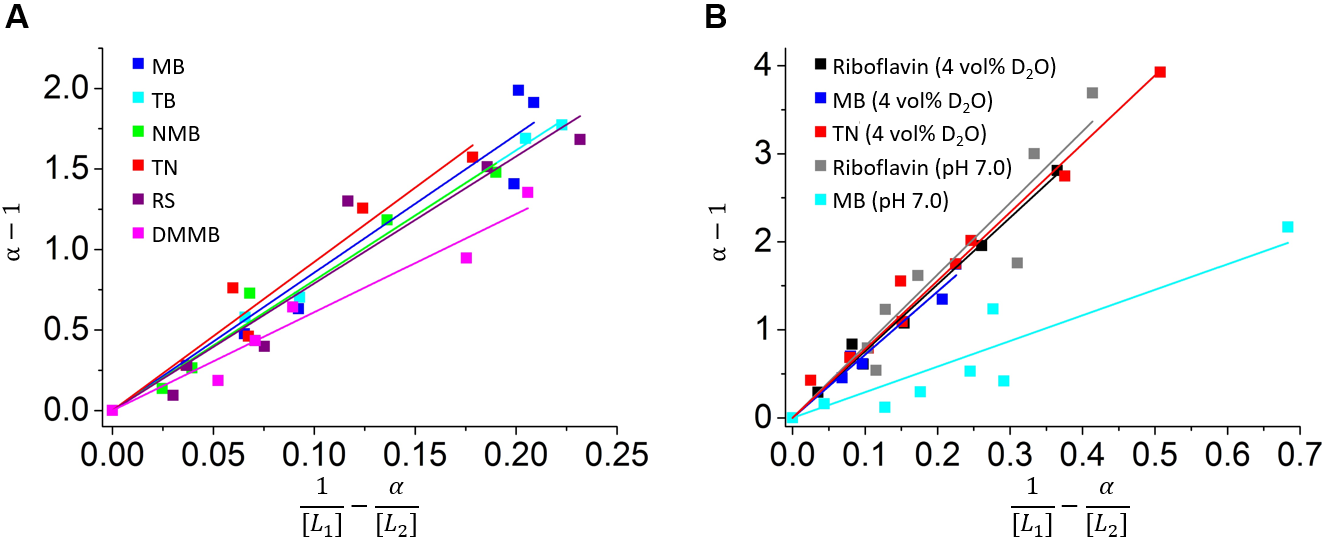

**Figure S9.** Estimation of dissociation constant (*K*_d_) about complex formation between Cyts and N5 molecules. *K*_d_ values were estimated via linear regressions, setting $\frac{1}{\left[ L \right]_{1}}-\frac{\alpha}{\left[ L \right]_{2}}$ and $\alpha-1$ as horizontal and vertical axis, respectively as described in eq. S2. (**A**) Plots of $\alpha-1$ against $\frac{1}{\left[ L \right]_{1}}-\frac{\alpha}{\left[ L \right]_{2}}$ at pH 7.8. The linear regressions are as follows; MB: b = 8.560a, R^2^ = 0.939; TB: b = 8.079a, R^2^ = 0.997; NMB: b = 8.071a, R^2^ = 0.949; TN: b = 9.227a, R^2^ = 0.949; RS: b = 7.890a, R^2^ = 0.927; DMMB: b = 6.013a, R^2^ = 0.958. (**B**) Plots of $\alpha-1$ against$\frac{1}{\left[ L \right]_{1}}-\frac{\alpha}{\left[ L \right]_{2}}$ at pH 7.0 or those in the presence of 4% (v/v) D_2_O at pH 7.8. The linear regressions are as follows; riboflavin with 4 vol% D_2_O: b = 7.579a, R^2^ = 0.987; MB with 4 vol% D_2_O: b = 7.172a, R^2^ = 0.972; TN with 4 vol% D_2_O: b = 7.781a, R^2^ = 0.977; riboflavin at pH 7.0: b = 8.137a, R^2^ = 0.906; MB at pH 7.0; b = 2.910a, R^2^ = 0.854. Numeric data of *K*_d_ are summarized in Table S1.

**A**

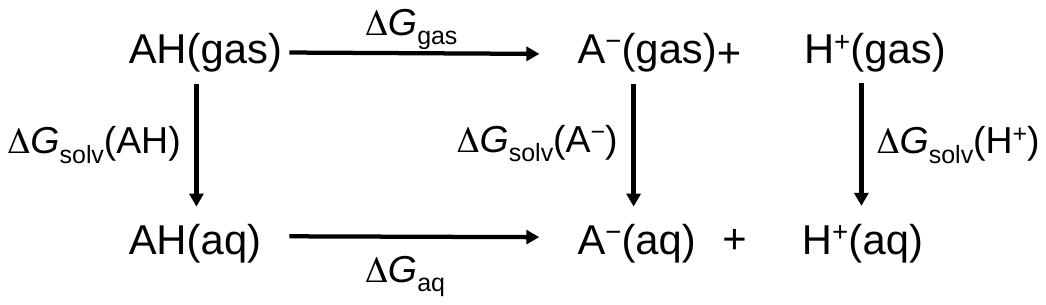

**B**

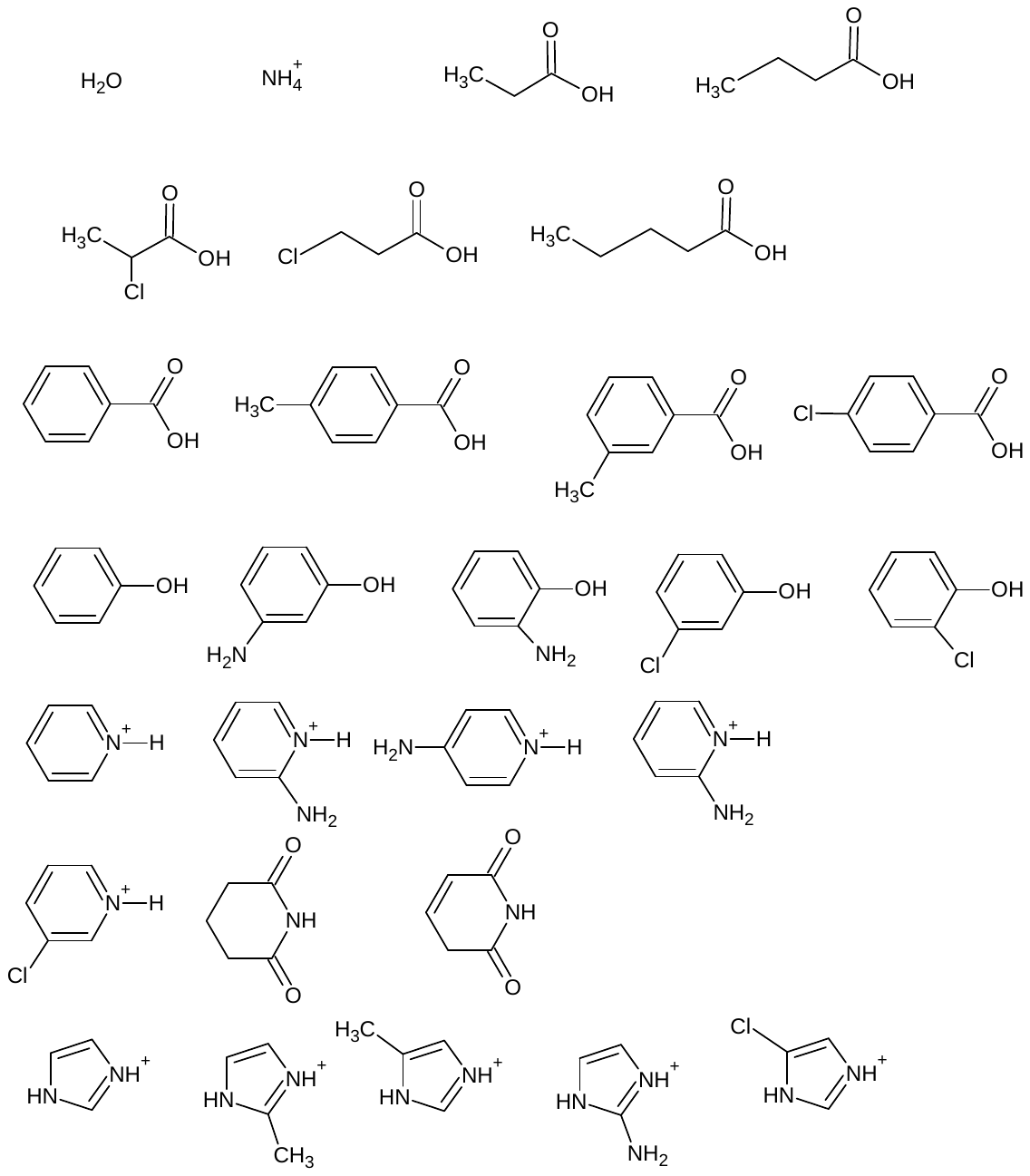

**Figure S10.** (**A**) Thermodynamic cycle for obtaining *ΔG*_aq_, according to eqs. 2 and 3. (**B**) 28 compounds evaluated in Figure. S11.

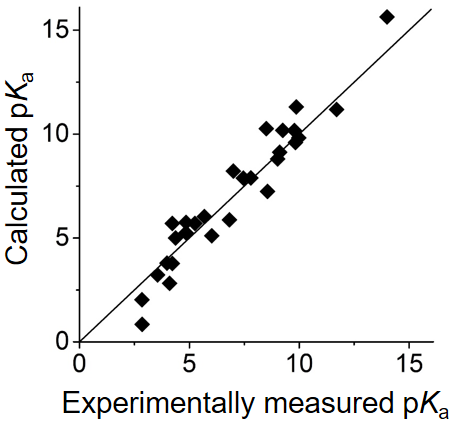

**Figure S11.** Correlation between experimentally measured and calculated p*K*_a_ values of 28 compounds in Figure S10. The root-mean-square deviation and the maximum error were 0.94 and 1.77 in p*K*_a_ units, respectively.

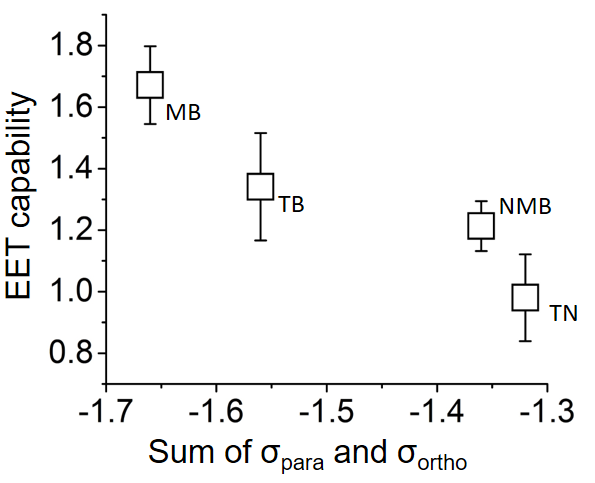

**Figure S12.** EET capabilities of *S. oneidensis* MR-1 with each N5 molecule as a function of the nucleophilicity at N5 site. Since MB, TB, NMB, and TN have the same heterocyclic backbone, the nucleophilicity of each molecule at N5 can be compared based on Hammett substituent constant. The nucleophilicity is calculated by the sum of the contribution from substituents located at para-position (σ_para_) and ortho-position (σ_ortho_). The molecule with more negative σ has higher nucleophilicity at N5. The numeric data of σ_para_ and σ_ortho_ are provided in Table S2.

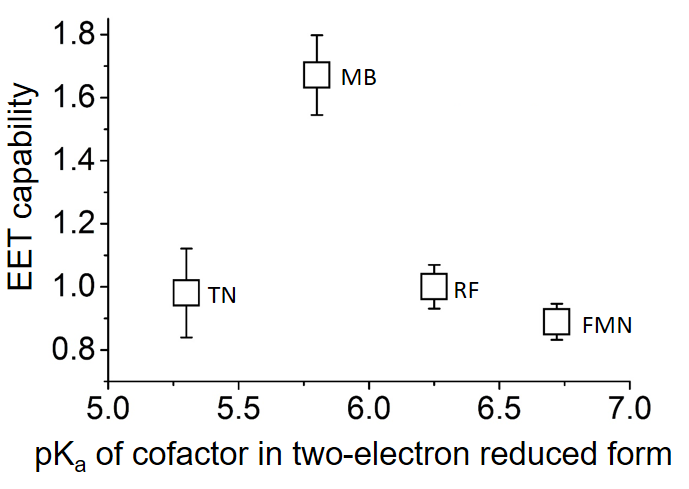

**Figure S13.** EET capabilities of *S. oneidensis* MR-1 with each N5 molecule as a function of the p*K*_a_ of cofactor in two-electron reduced form. The p*K*_a_ of each cofactor was obtained by refs. ^4-6^.

**Table. S1 Quantification of EET capability with each redox molecule normalized by the amount of cofactor-bound Cyts**

|  | Maximum current production with 2.0 μM molecule (*I_c_*) /μA | Dissociation constant (*K*_d_) /μM | EET capability |
| --- | --- | --- | --- |
| FMN | 26.73±1.72 | 10^*^ | 0.89±0.06 |
| Riboflavin | 30.07±2.08 | 10^*^ | 1.00±0.07 |
| MB | 57.15±3.12 | 8.56±0.55 | 1.67±0.13 |
| TB | 48.03±6.22 | 8.08±0.12 | 1.34±0.17 |
| NMB | 43.47±2.32 | 8.07±0.41 | 1.21±0.08 |
| TN | 31.51±4.24 | 9.23±0.57 | 0.98±0.14 |
| RS | 28.08±3.30 | 7.89±0.59 | 0.77±0.10 |
| DMMB | 25.78±1.46 | 6.01±0.34 | 0.57±0.04 |

*Values from ref. ^7-8^.

**Table. S2 Hammett parameters (σ_para_ and σ_ortho_) and the sum of σ_para_ and σ_ortho_ (Σσ) to N5 site of MB, TB, NMB, and TN** ^9-10^

|  | σ_para_ | σ_ortho_ | Σσ |
| --- | --- | --- | --- |
| MB | -1.66 | 0 | -1.66 |
| TB | -1.49 | -0.07 | -1.56 |
| NMB | -1.22 | -0.14 | -1.36 |
| TN | -1.32 | 0 | -1.32 |

σ_para_ for NH_2_: -0.66; σ_para_ for N(CH_3_)_2_: -0.83; σ_para_ for NH(C_2_H_5_): -0.61; σ_ortho_ for CH_3_: -0.07
